# If it’s real, could it be an eel?

**DOI:** 10.1101/2023.01.07.523085

**Authors:** Floe Foxon

**Affiliations:** Department of Data Management and Statistical Analysis, PinneyAssociates – Pittsburgh, USA

## Abstract

Previous studies have estimated the size, mass, and population of hypothetical unknown animals in a large, oligotrophic freshwater loch in Scotland based on biomass and other observational considerations. The ‘eel hypothesis’ proposes that the anthrozoological phenomenon at Loch Ness can be explained in part by observations of large specimens of European eel (*Anguilla anguilla*), as these animals are most compatible with morphological, behavioural, and environmental considerations. The present study expands upon the ‘eel hypothesis’ and related literature by estimating the probability of observing eels at least as large as have been proposed, using catch data from Loch Ness and other freshwater bodies in Europe. Skew normal and generalized extreme value distributions were fitted to eel body length distributions in order to estimate cumulative distribution functions from which probabilities were obtained. The chances of finding a large eel in Loch Ness are around 1 in 50, 000 for a 1-meter specimen, which is reasonable given the loch’s fish stock and suggests some sightings of smaller ‘unknown’ animals may be accounted for by large eels. However, the probability of finding a specimen upwards of 6 meters is essentially zero, therefore eels probably do not account for ‘sightings’ of larger animals. The existence of exceedingly large eels in the loch is not likely based on purely statistical considerations.

## Introduction

Loch Ness is a large, oligotrophic freshwater loch located along the Great Glen Fault in Scotland. Since the 1930s, purported sightings of unknown animals in the loch have featured prominently in popular media, though to date no specimen has been obtained despite numerous efforts, making the probability of such animals unlikely.

The authenticity and interpretations of photographs and films allegedly depicting unknown animals in Loch Ness have been seriously doubted (Martin & Boyd 1999; Shine 2003; Raynor 2010; Campbell 1986; Raynor 2009). In the 20^th^ century, systematic searches with submersibles, sector-scanning sonar surveys, hydrophones, underwater photography, longlining, and trawling undertaken by the Loch Ness Investigation Bureau (LNIB) (Mackal 1976, Chapters II–V), the Academy of Applied Science (AAS) (Scott & Rines 1975; Klein et al. 1972; Rines 1982; Rines & Curtis 1979; Rines et al. 1984; Edgerton et al. 1989), and the Loch Ness and Morar Project (LNMP) (Shine 2006) have returned only ambigious sonar signals, low-quality photographs, and unidentifiable sound recordings.

In the 1970s, a sample of European eels (*Anguilla anguilla*) were collected from Loch Ness with baited traps. The distribution of eel masses was skewed, which led biologist Roy Mackal (1976) to conclude that large eels may exist in the loch. Eel body structure and function are characterised by an elongated body form, a single pair of pectoral fins, strong musculature and high-amplitude winding movement, and durable integument with thick epidermis and dark chromatophores (Kloppmann 2003).

Mackal noted that large eels would therefore be consistent with eyewitness descriptions of Loch Ness animals including reports of an elongated head-neck, pectoral fins, extreme flexion, and dark integument (Gibson & Heppell 1988a,b). This explanation has also been reviewed by naturalists Adrian Shine and David Martin of the LNMP, who noted that eels migrate via the River Ness and that most ‘sightings’ occur near river mouths.

An eDNA sudy conducted at the loch in 2018 detected extraordinary amounts of mtDNA and nuDNA from eels (Shine 2019), prompting principal investigator Neil Gemmell to further suggest the possibility of large eels in the loch (University of Otago 2019). A large eel-shaped animal was recently filmed in the River Ness by the Ness Fishery Board (Turner 2019).

Simon (2007) suggests the physiologically possible maximum length of *A. anguilla* is 0.5–1.3 meters, which is not particularly monstrous. Using wave mechanics, LeBlond & Collins (1987) estimate the size of the subject depicted in the infamous “Surgeon’s Photograph” at Loch Ness at 0.6–2.4 meters.

Much larger estimates have been made. Based on their “flipper” photograph, Scott & Rines (1975) estimated a total body length of 15–20 meters for an unknown Loch Ness animal. These estimates seem inconsistent with Loch Ness biomass calculations. Sheldon & Kerr (1972, 1973) estimate a population of 156 if the individual mass of a hypothetical unknown animal in Loch Ness is 100 kilograms, and suggest just one individual can exist if its mass is 2,000–3,000 kilograms. Scheider & Wallis (1973) estimate the loch may support 157 animals of 100 kilograms each, and 10 animals of 1,500 kilograms each.

Finally, in an article titled “If there are any, could there be many?”, Carl Sagan (1976) used collision physics and suggested that a population of 300 animals each of 10-meter size would be consistent with AAS observations. Thus, if there are any, there may be many. If it’s real, could it be an eel? The aim of the present study is to estimate the probability of finding various sizes of eel in Loch Ness based on available catch data.

## Methods

Data on the mass distribution of *A. anguilla* in Loch Ness were taken from *n* = 129 eels caught between 1970 and 1971 under the supervision of the LNIB and described in Mackal (1976, Appendix G).

Mackal does not provide the length distribution for the sample, nor did Mackal directly regress length on mass. Mackal did regress functions of length on mass and maximum circumference, but provided only non-linear equations unsuitable for solving simultaneously. Thus, to convert the mass distribution into a length distribution, the relationship between mass and length in *A. anguilla* captured in Scottish freshwaters from 1986–2008 was taken from Figure 2 of Oliver et al. (2015) as

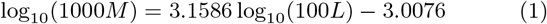

for mass *M* in kilograms and length *L* in meters. This equation is used over alternatives such as that in Figure 2(b) of Simon (2007) because Equation 1 is most relevant to Loch Ness (having been derived from Scottish freshwater).

Rearranging Equation 1 gives

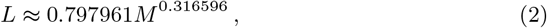

from which the lengths of eels in Mackal’s sample were estimated.

Skew normal and generalized extreme value distributions were fitted to the Loch Ness eel length distribution, and fit parameters were used to estimate the probability density function (PDF) and cumulative distribution function (CDF).

The probability of finding an eel in Loch Ness at least as long as *L* was then estimated from the CDF as>

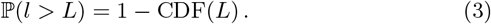

For comparison, the above analysis was repeated with publicly-available length data on *n* = 420 European eels captured in Zeeschelde, Belgium presented in Verhelst et al. (2018, Supplemental Information). Zeeschelde was selected because raw data were available, whereas this is not the case for other published analyses.

All analyses were performed in Python 3.8.8 with the packages Numpy 1.20.1, Pandas 1.2.4, Matplotlib 3.3.4, and Scipy 1.6.2. All code and data are available in the online Supplementary Information (Foxon 2022). Maps of locations referenced throughout the manuscript in order of reference are also available in the online Supplementary Information.

## Results

The length distributions for the Loch Ness and Zeeschelde eel samples are shown in the upper plots of Figure 1. Fits were similar for both skew normal and generalized extreme value distributions. The distribution for Loch Ness is evidently skewed toward lower lengths, similar to the length distribution of European eels presented in Fig. 2 of Melia et al. (2006) for eels captured in the Camargue lagoons of France. The Zeeschelde distribution has notably less skew. The length distribution in a sample of eels in Vistonis Lake, Greece was skewed toward higher lengths (see Fig .2 of Macnamara et al. (2014)), suggesting much variability in eel length skew across different environments.

**Figure 1.**
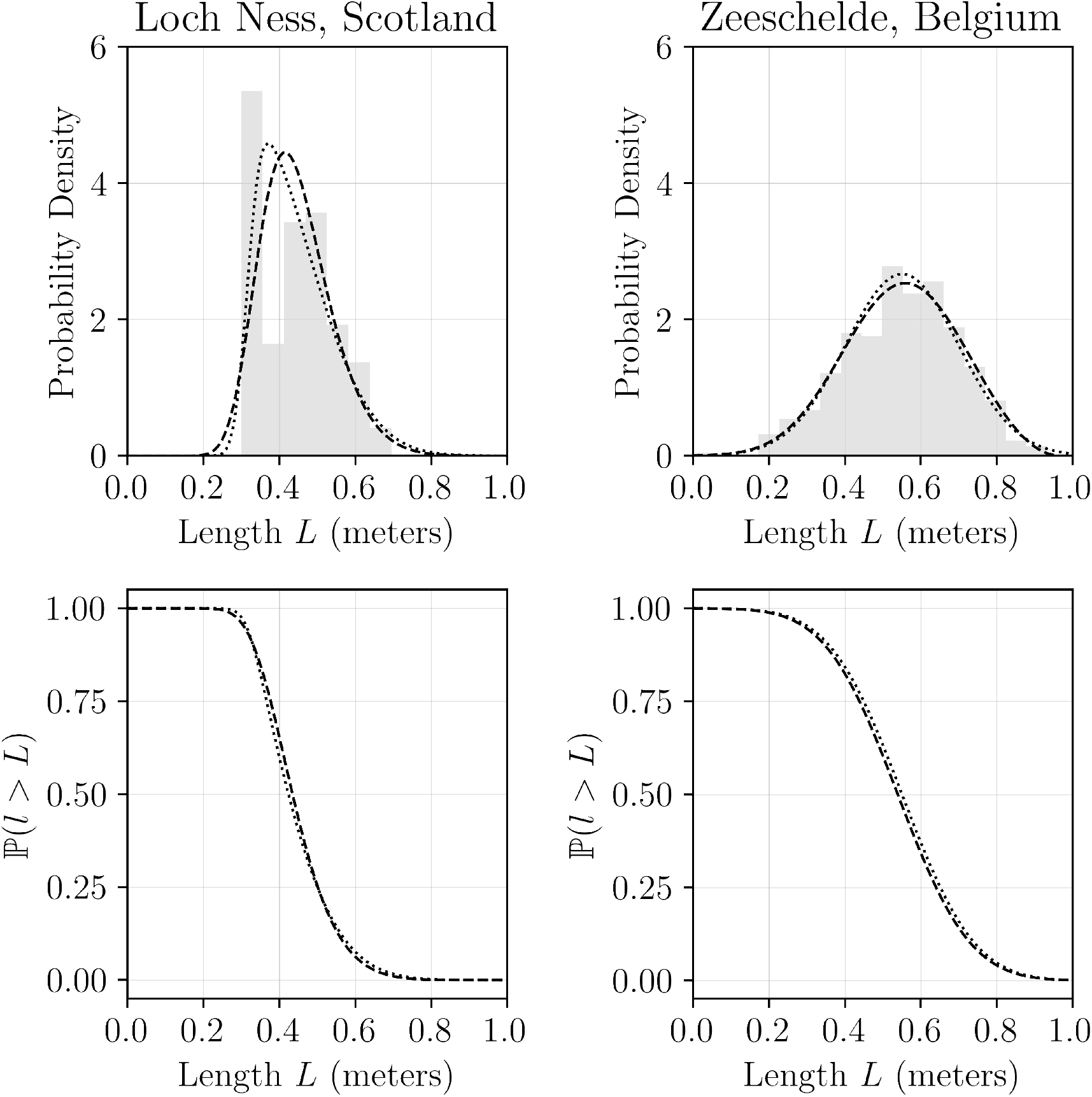
Length distributions for European eels (*A. anguilla*) captured in Loch Ness (left) and in Zeeschelde (right). The lower plots show the probability of finding an eel at least as long as *L*. Dotted lines represent skew normal distribution fits. Dashed lines represent generalized extreme value distribution fits. The number of bins follows the rule provided in Freedman & Diaconis (1981).

**Figure 2.**
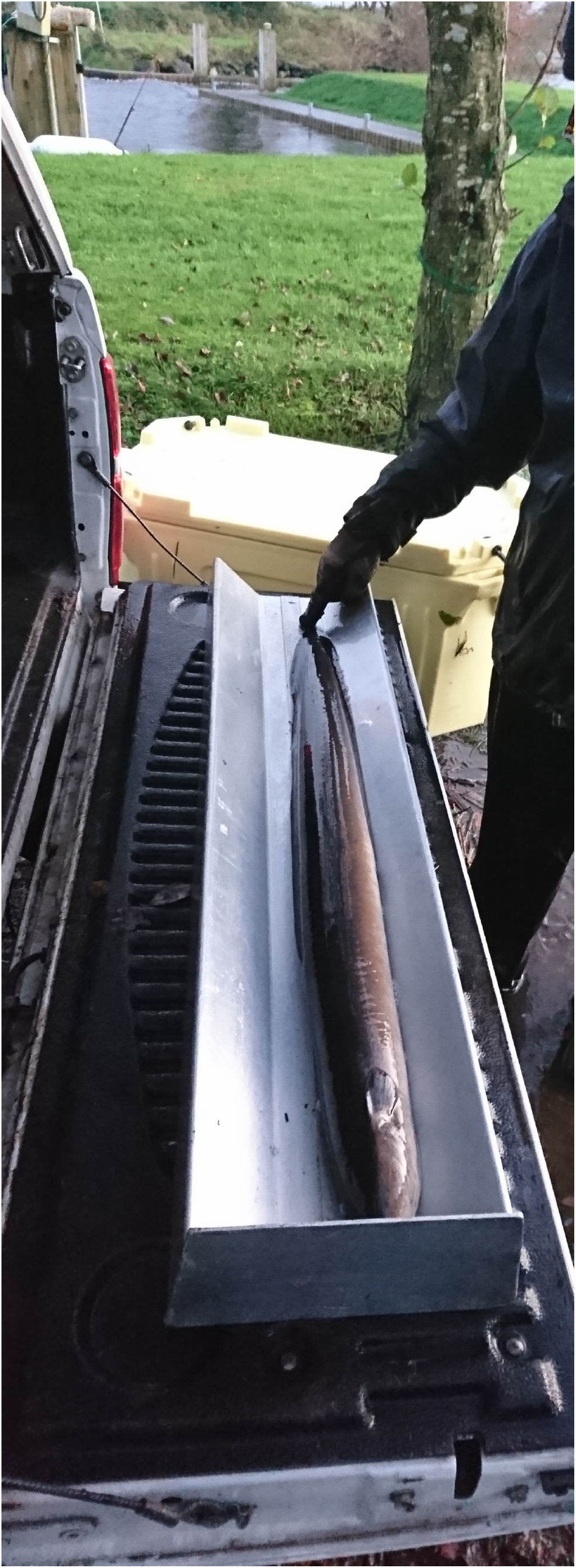
1.05-meter female silver eel. Image courtesy of Dr Derek W Evans, AFBINI.

The probabilities of finding an eel at least as big as *L* in Loch Ness and in Zeeschelde are shown in the lower plots of Figure 1. Loch Ness has a comparatively low probability of finding larger eels 0.6–0.8 meters in length, and the probability of finding an eel in excess of 1 meter length is very low for both environments.

Table 1 contains the estimated probabilities for specific lengths of interest. Results were somewhat similar for skew normal and generalized extreme value fits, and for both Loch Ness and for Zeeschelde.

**Table 1.**
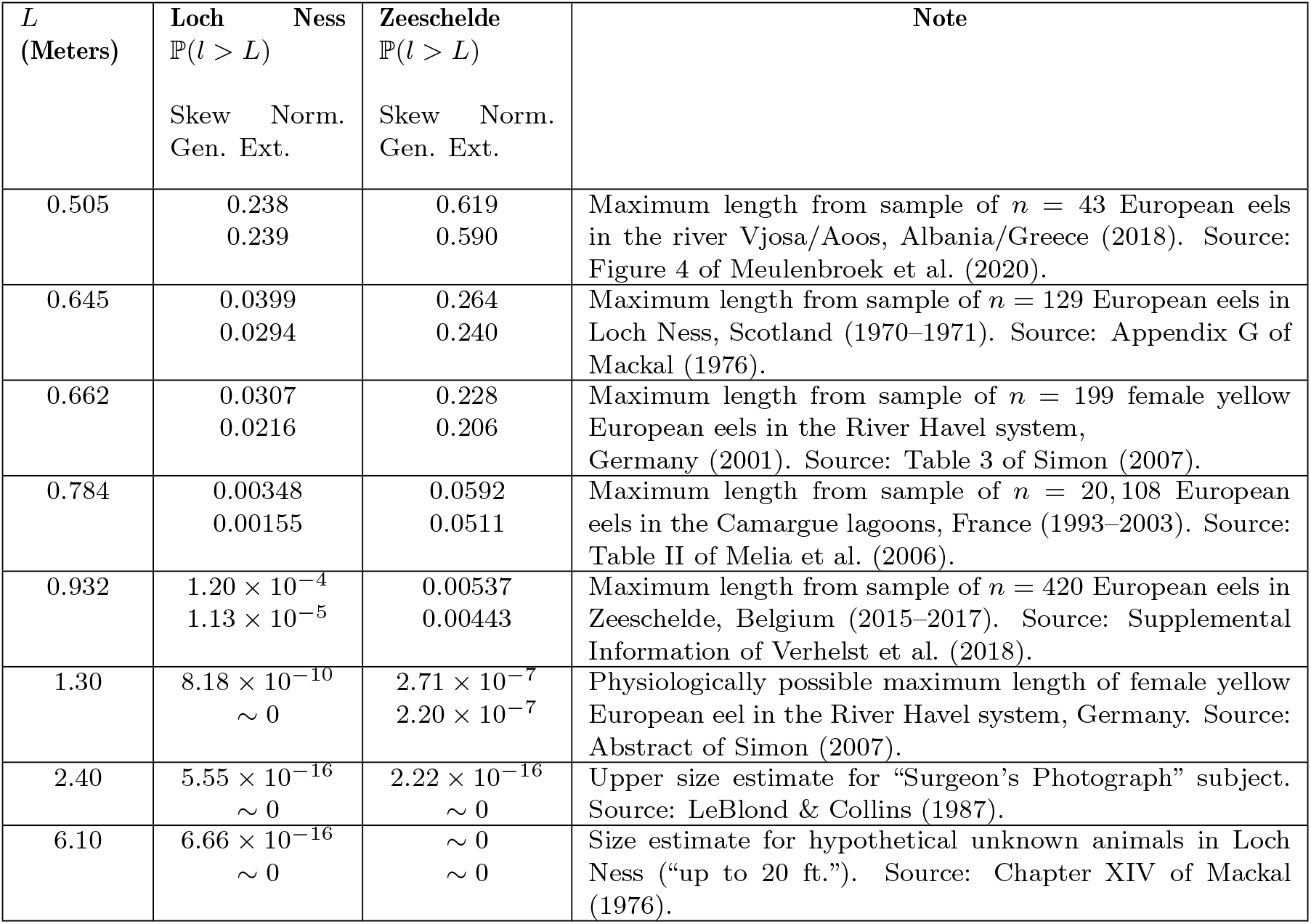
Probabilities associated with finding a European eel (*A. anguilla*) at least as large as length *L* in Loch Ness and Zeeschelde for various *L*.

While the chance of finding a large eel approximately 1 meter in length in Loch Ness is low (around 1 in 50, 000), this is certainly possible given the eel population of the loch: assuming a standing fish stock of 0.55 kg ha^*−*1^ for Loch Ness (Sheldon & Kerr 1972, 1973) and given a surface area of 5, 600 hectares, the total standing fish stock of the loch is approximately 3, 080 kg. Further assuming that half of this stock is eel mass (plausible given the Loch Ness eDNA study (Shine 2019)), this would imply 1, 540 kg of eel in the loch. The average Loch Ness eel mass is 0.1857 kg (Mackal 1976, Appendix G), therefore there are over 8, 000 eels in Loch Ness at a given time. Over the course of a few generations, an eel 1 meter in length may be expected.

However, this is not quite the ‘monster’ postulated. Indeed, the probability of finding a 6-meter eel in Loch Ness is essentially zero; too low for the software used to provide a reliable estimate. Thus, while large eels may account for some eyewitness sightings of large, animate objects rising to the loch surface, they are unlikely to account for ‘sightings’ of extraordinarily large animals, which may instead be accounted for by wave phenomena, the occasional stray mammal, or other.

## Discussion

This study used data on the distribution of European eel (*A. anguilla*) masses in an oligotrophic freshwater loch in Scotland to estimate the probability of finding an eel of extraordinary size there. Similar to other eel populations in Europe, the average eel length in Loch Ness is relatively small, and highly comparable to biometric data on silver eels (sexually mature *A. anguilla*) collected in the same decade at the Girnock Burn fish trap on the River Dee in Aberdeenshire, Scotland (see Table 5.3 of Bašić et al. (2021)), suggesting Loch Ness eels are similar to those found elesewhere in the country. Findings of the present study suggest that the chance of finding a 1-meter eel in the loch (1 in 50, 000) is reasonable given the standing fish stock, and so some eels may account for purported sightings of somewhat large animals at the loch surface. For comparative purposes, Figure 2 shows an image of a female silver eel 1.05 meters in length. Other images of European eels in excess of 1-meter in length are provided in the online Supplementary Information (Foxon 2022).

However, the present analyses suggest that larger eels upwards of 6 meters are highly improbable, therefore ‘super’ eels are an unlikely explanation for eyewitness reports of the very largest alleged animals at Loch Ness. Marine Scotland Science have reported growth rates of eels on the Girnock tributary of the River Dee in Scotland as high as 35.2 mm yr^*−*1^ (Bašić et al. 2021). Assuming a linear rate of growth throughout the lifecyle of an eel (in reality the rate of change of an eel’s total length is an exponential decay curve, as on page 175 of Tesch & Thorpe (2003b)), it would take an eel in Scotland almost 30 years at such a growth rate to reach the 1-meter size. Given that the oldest eels can live for a number of decades (Tesch & Thorpe 2003b), again it seems likely that approximately 1 meter is a realistic maximum length for eels in Loch Ness; a 6-meter specimen would need to live with constant high growth for almost 200 years, an age approaching that of the longest-living fish, the Greenland shark *Somniosus microcephalus*. Though one

European eel reportedly (unverified) lived to the grand age of 155 years (Rundquist 2014), that specimen did not grow to a remarkable size, again because eel growth is non-linear, slowing in older ages. Furthermore, the “breaching” behaviour attributed to unknown Loch Ness animals (swimming upwards and out of the water) is not a behaviour that is characteristic of eels during migration or otherwise (Tesch & Thorpe 2003b), especially as such behaviour would represent unnecessary energy expenditure in a cold environment with relatively little food.

This analysis is limited by several factors. First, the Loch Ness eel sample used was relatively small at *n* = 129. Larger samples across longer time periods may provide more accurate estimates. Second, the assumption of a skew normal distribution would not hold if, for example, a larger sample revealed a bimodal distribution of eel lengths with a small peak at higher lengths. Third, this analysis is based on purely statistical considerations; the biological mechanism behind the physiological possibility of much larger eels is beyond the scope of the present study. Some authors have suggested one such mechanism as neoteny (Mackal 1976, Chapter XI), i.e. uncontrolled growth of the leptocephalus larva in *A. anguilla* preceding subsequent stages of development (Tesch & Thorpe 2003a). Fourth, environmental conditions such as temperature and available biomass impact eel growth and length, therefore comparisons to other environments such as Zeeschelde may not be appropriate, i.e., some of the data cited may not be relevant to the relatively cold waters of Loch Ness.

In conclusion, while Sagan (1976) found that if there are any, there may be many, the present study shows that if it’s real, it could be an eel, but not a very large one…

## Acknowledgements

The author thanks Charles G.M. Paxton, Dr Don Jellyman, and Dr Derek W Evans for helpful comments and suggestions.

## Conflict of interest disclosure

The author declares that they have no financial conflicts of interest in relation to the content of the article.

Data, script, code, and supplementary information availability

## Data and code are available online

https://doi.org/10.17605/OSF.IO/ZMGDJ

